# Insulin signaling couples growth and early maturation to cholesterol intake

**DOI:** 10.1101/2021.06.07.447368

**Authors:** Michael J. Texada, Mette Lassen, Lisa H. Pedersen, Alina Malita, Kim Rewitz

## Abstract

Nutrition is one of the most important influences on growth and the timing of developmental maturation transitions including mammalian puberty and insect metamorphosis. Childhood obesity is associated with precocious puberty, but the assessment mechanism that links body fat to early maturation is unknown. During development, intake of nutrients promotes signaling through insulin-like systems that govern the growth of cells and tissues and also regulates the timely production of the steroid hormones that initiate the juvenile-adult transition. We show here that the dietary lipid cholesterol, required as a component of cell membranes and as a substrate for steroid biosynthesis, also governs body growth and maturation in *Drosophila* via promoting the expression and release of insulin-like peptides. This nutritional input acts via the Niemann-Pick-type-C (Npc) cholesterol sensors/transporters in the glia of the blood-brain barrier and cells of the adipose tissue to remotely drive systemic insulin signaling and body growth. Furthermore, increasing intracellular cholesterol levels in the steroid-producing prothoracic gland strongly promotes endoreduplication, leading to accelerated attainment of a nutritional checkpoint that ensures that animals do not initiate maturation prematurely. These findings couple sensing of the lipid cholesterol to growth control and maturational timing, which may help explain both the link between cholesterol and cancer as well as the critical connection between body fat (obesity) and early puberty.

**HIGHLIGHTS:**

- Dietary cholesterol promotes developmental growth and leads to early maturation
- Insulin signaling couples cholesterol intake with systemic growth
- Cholesterol promotes insulin signaling and growth via glial and fat-tissue relays
- Cholesterol sensing affects a nutritional checkpoint that prevents early maturation

GRAPHICAL ABSTRACT

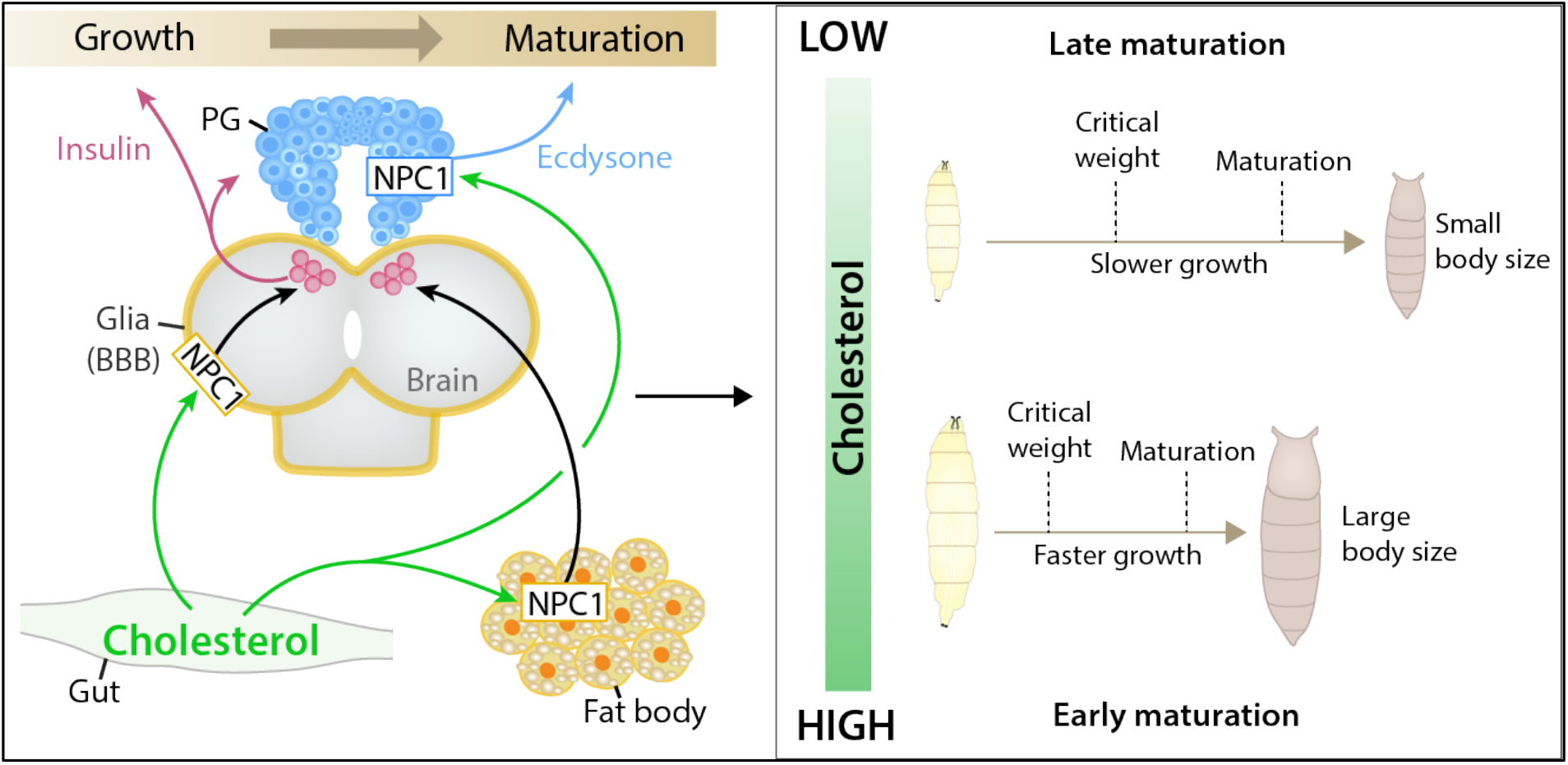

## INTRODUCTION

Animals’ growth and development depend upon nutrient availability, and therefore specialized cells and tissues have arisen that sense nutritional inputs and adjust growth and developmental programs via systemic hormonal pathways [1]. In most eumetazoans, these include the conserved insulin-like-peptide and steroid-hormone signaling systems. These become dysfunctional when nutrient levels exceed their evolutionarily normal range. Overloading of the insulin system leads to obesity, metabolic syndrome, insulin resistance, and other pathophysiologies, and overnutrition also leads to precocious puberty associated with childhood obesity [2].

Many animals’ early life is a non-reproductive stage of rapid growth, terminated at some nutritional threshold that signals the animal’s readiness to become a reproductively fit adult [1]. In animals as diverse as humans and insects, this transition is driven by steroid hormones [3] – gonadal steroids including testosterone and estrogen trigger mammalian puberty, and insect metamorphosis is initiated by ecdysone, produced in the prothoracic gland (PG). Similar neuroendocrine cascades regulate insect and mammal steroidogenesis, including the orthologous neuropeptides Allatostatin A/Kisspeptin and Corazonin/gonadotropin-releasing hormone (GnRH) [4, 5] as well as analogous steroid-feedback circuits [6, 7].

These axes are clearly linked to the metabolic state of the animal, including attainment of a certain critical size [1]. However, the mechanisms of body-size estimation and the effects of nutritional status are not completely understood. Recent work in *Drosophila* suggests that progression to adulthood is gated by a checkpoint system monitoring tissue growth and nutritional status [8]. When animals reach a “critical weight” (CW), they become committed to completing their development and maturation irrespective of further nutritional inputs, whereas animals starved before this checkpoint is satisfied halt their progression to adulthood. This suggests that the CW checkpoint assesses the animal’s nutritional state, but the specific nutrients required, and their levels are sensed, are incompletely understood.

In *Drosophila*, nutritional input drives growth and maturation through the insulin pathway. Nutrient intake, particularly of amino acids, is sensed via the fat body (analogous to mammalian adipose tissue) and the glia of the blood-brain barrier (BBB). These tissues release factors that regulate the expression and release of *Drosophila* insulin-like peptides (ILPs) from the insulin-producing cells (IPCs) within the brain [9–11], which share functional and developmental homology with mammalian pancreatic beta cells [1]. These ILPs promote systemic growth through the conserved insulin-receptor/PI3K/Akt pathway. Insulin also promotes PG ecdysone production, linking nutrition directly to developmental progression [12–16].

Human puberty-triggering “critical weight” appears linked to body-fat stores [17], which may explain the link between childhood obesity and early puberty. Despite this, the mechanism by which adiposity affects puberty initiation is unclear; furthermore, the role of cholesterol has not been considered, even though adipose tissue is a major cholesterol storage depot, especially in obesity. Sterols such as cholesterol have membrane-structural functions but also play important signaling roles [18, 19], and sterols are required as substrates for steroid-hormone production. Insects, including *Drosophila*, have lost the ability to synthesize sterols *de novo* and thus must acquire them through feeding [20]. Mammals are cholesterol prototrophs, but most intracellular cholesterol still comes from low-density-lipoprotein (LDL)-mediated cellular uptake of dietary cholesterol [21]. In both taxa, consumed sterols are transported in the circulatory system bound within lipoprotein particles (“LPPs” such as mammalian LDL/HDL), and target tissues take them up through a variety of mechanisms including receptor-mediated endocytosis [22]. LPP-bound sterols are extracted in the lysosome and inserted into the lysosomal membrane by membrane-integral transport proteins. The primary such protein, Npc1, underlies the Niemann-Pick type C lysosomal storage disorder [23]; without Npc1 function, cholesterol accumulates in endosomal-lysosomal compartments, leading to increased intracellular cholesterol signaling. Thus, Npc1 seems to be part of a mechanism by which cells sense cholesterol [24].

We wished to determine the routes by which cholesterol regulates *Drosophila* larval growth. Our findings show that dietary cholesterol dose-responsively promotes growth and accelerates development by enhancing insulin signaling; cholesterol sensing in the cells of the fat body and the glia of the BBB remotely induces the release of ILPs from the IPCs. Manipulating cholesterol sensing in the PG also promotes endoreduplication and leads to premature attainment of the CW checkpoint. Thus, dietary cholesterol accelerates growth through insulin signaling and leads to early maturation through effects on steroidogenesis.

## RESULTS

### Dietary cholesterol promotes systemic growth via insulin signaling

Growth occurs almost exclusively during larval life in *Drosophila*; thus pupariation fixes animal size. We reared larvae on synthetic diets [25] containing a range of cholesterol concentrations and measured their pupariation timing and pupal size. We found that larvae fed higher-cholesterol diets pupariated much earlier than animals reared on lower concentrations but nevertheless formed larger pupae (Fig. 1A and 1B), indicating an increased growth rate during their shortened growth period. Thus, the intake of this nutrient is somehow coupled to systemic growth and development.

**FIGURE 1.**
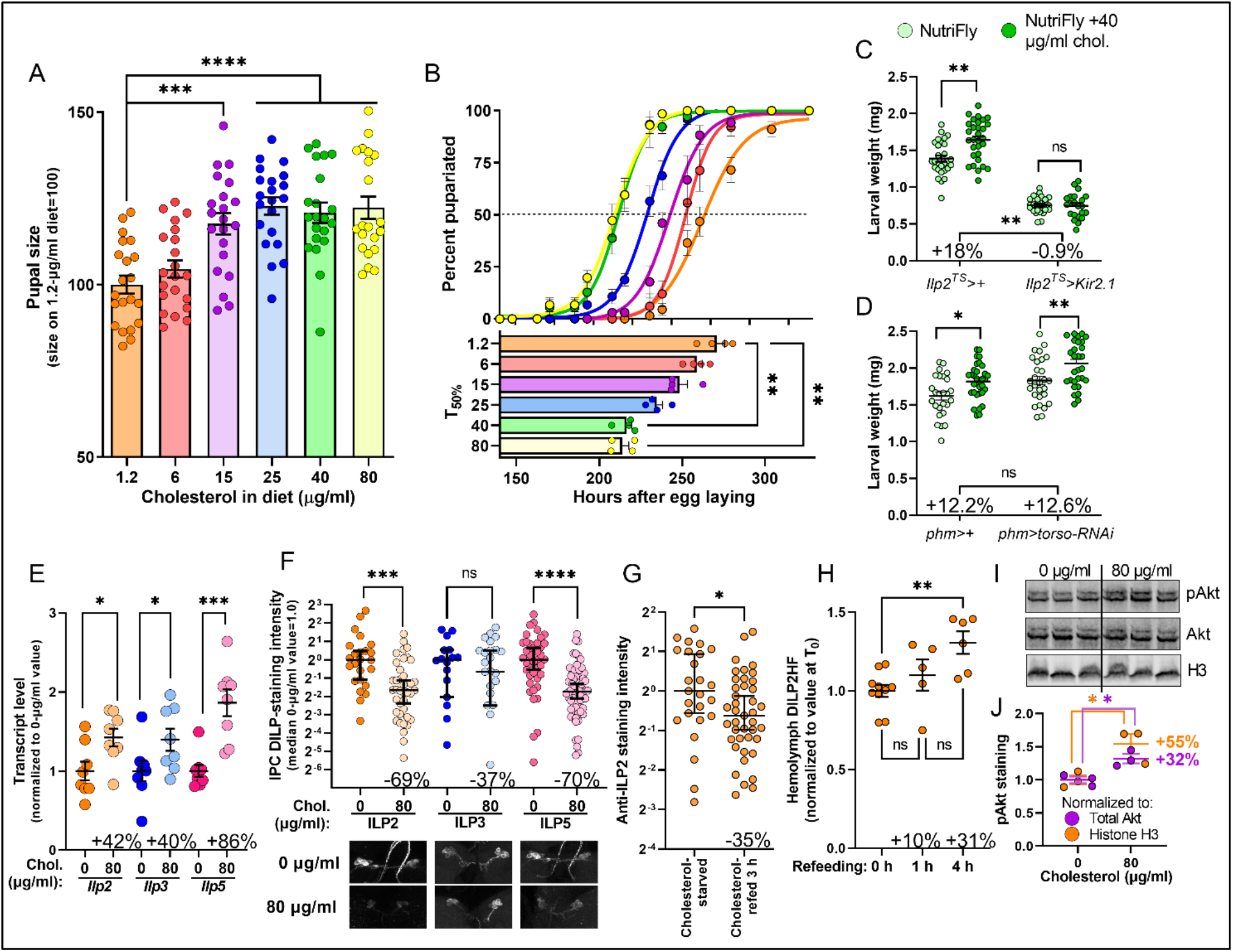
Dietary cholesterol systemically promotes growth through insulin signaling. **A:** Larval growth on synthetic medium, reflected in pupal size, increases with increasing dietary cholesterol concentration. **B:** Development to the pupal stage on synthetic medium is accelerated with increasing dietary cholesterol concentration. Top: fraction pupariated over time; bottom: time until 50% pupariation; each data point represents a vial of flies. **C:** Cholesterol’s growth-promoting effect requires insulin signaling. Dietary supplementation with cholesterol (NF vs. NF+40) promotes growth in control animals, but if the IPCs are silenced through expression of the inhibitory channel Kir2.1 (*Ilp*2^ts^>*Kir2.1*), this growth promotion is blocked (2-way ANOVA *p* for interaction of genotype and diet, 0.0105). **D:** The growth-promoting effect of added cholesterol does not arise through effects on ecdysone production. Animals were reared on NF medium until 72 hours AEL, when half were transferred to NF+40. Larvae were weighed at 104 hours AEL. Blocking the production of ecdysone by expressing RNAi against the PTTH receptor Torso in the prothoracic gland had no effect on cholesterol-induced growth (2-way ANOVA *p* for interaction of genotype and diet, 0.974). See also Fig. S1. **E and F:** Insulin-gene expression is higher (**E**), and anti-DILP staining intensity is lower (**F**), in animals fed medium containing 80 μg/ml cholesterol than in animals fed cholesterol-free medium, suggesting increased DILP on cholesterol-containing medium. Illustrative images are shown below. **G:** Cholesterol refeeding can acutely induce DILP2 release within 3 hours, as indicated by reduced anti-DILP2 stain in the IPCs. **H:** Hemolymph ELISA against tagged DILP2 indicates correspondingly increased circulating DILP2 after larvae are transferred from cholesterol-free diet to cholesterol-replete medium. **I, J:** Peripheral insulin-signaling activity is increased by cholesterol refeeding, reflected in anti-phospho-Akt staining normalized to total Akt or to histone H3. Statistics: A: mean±SEM; Welch’s ANOVA with Dunnett’s T3 multiple comparisons. B: top, mean±SEM; bottom, mean±SEM with Kruskal-Wallis ANOVA with Dunnett’s multiple comparisons. C and D, mean±SEM, with two-way ANOVA with Šídák’s multiple comparisons. E: mean±SEM with Mann-Whitney pairwise test. F: median with 95% confidence interval, shown on log scale, with Mann Whitney pairwise comparisons. G: median with 95% confidence interval, shown on log scale, with Mann Whitney pairwise comparison. H: mean±SEM; Kruskal-Wallis ANOVA with Dunn’s multiple comparisons. J: mean±SEM with unpaired t-test. Significance is noted as *, *p*<0.05; **, *p*<0.01; ***, *p*<0.001; ****, *p*<0.0001.

We next wished to determine the basis for this effect. Systemic growth is largely governed by the antagonistic insulin and ecdysone signaling pathways [1]. We therefore asked whether either of these mediates the effects of dietary cholesterol on growth. The ILPs central to systemic growth regulation are ILP2, ILP3, and ILP5, produced by the IPCs [26, 27]. These peptides promote body growth, and thus increased ILP activity shortens the time required to reach critical weight [28]. The effects observed above are consistent with increased insulin-signaling activity, so we assessed the necessity of this pathway for cholesterol-induced growth enhancement. Completely synthetic diet is sub-optimal, as reflected in the prolonged developmental times above; therefore we used a commercial low-sterol cornmeal medium (“NutriFly”, NF) that can be supplemented with cholesterol to promote growth [29]. We inhibited ILP secretion in the early third-instar stage using *Ilp2-GAL4* together with *Tubulin* (*Tub*)-driven temperature-sensitive GAL80 (together, “*Ilp2^TS^*>”) to drive expression of the inhibitory potassium channel Kir2.1 in the IPCs. Control animals (*Ilp2^TS^*>+) grew significantly more quickly after transfer to higher-cholesterol diet, whereas increased dietary cholesterol had no effect on the larval weight of *Ilp2^TS^*>*Kir2.1* animals (Fig. 1C; 2-way ANOVA genotype×diet interaction *p*=0.0024). Thus, ILP release is required for the growth-enhancing effects of dietary cholesterol. Importantly, this also demonstrates that cholesterol’s growth-promoting effect does not merely reflect (for example) membrane-structural requirements.

Cholesterol is also the chemical precursor to the steroid hormone ecdysone, which has the effect of reducing the growth of larval tissues [15, 29]. Therefore cholesterol supplementation during larval life could alter growth through enhanced ecdysone production, although this would be expected to *decrease* systemic growth. To exclude the involvement of ecdysone, we reduced steroidogenesis through inhibition of PTTH-Torso signaling [30], using PG-specific *phantom-GAL4* (*phm*>) to drive RNAi against *torso*, thereby blocking PG activation. Unlike IPC-inhibited animals (Fig. 1C), PG-inhibited animals (*phm*>*torso-RNAi*) exhibited cholesterol-induced growth increase similar to controls’ (Fig. 1D, genotype×diet interaction *p*=0.76). Furthermore, we compared the growth of wild-type (*w*^1118^) animals fed synthetic diets containing either 1 or 80 μg/ml cholesterol, with and without direct supplementation with 250 μg/ml 20-hydroxyecdysone (20E), the active form of ecdysone. Added 20E attenuated larval growth on both diets, as expected, and cholesterol induced a similar size increase whether or not ecdysone was present (Fig. S1, 2-way ANOVA *p* for interaction, 0.65). Thus, cholesterol-induced systemic growth increase does not arise through altered ecdysone signaling but does require insulin signaling.

### Dietary cholesterol promotes the expression and release of ILP2 and ILP5

To assess the effects of cholesterol feeding on the insulin system, we measured the production and release of ILP2, −3, and −5 on otherwise-identical cholesterol-replete and -free diets. Expression of *Ilp2*, *Ilp3*, and *Ilp5* was significantly upregulated after 8 hours’ feeding on cholesterol-containing food (Fig. 1E). We next measured the amounts of ILP2, ILP3, and ILP5 present within the IPCs of these animals via immunostaining as an indicator of peptide release or retention. Despite the increased insulin expression observed in cholesterol-fed animals, these animals exhibited strongly reduced IPC ILP2 and ILP5 staining levels (Fig. 1F), suggesting enhanced release of these peptides. To test whether cholesterol can induce ILP release acutely, we cholesterol-starved animals for 16 hours and then transferred them to a cholesterol-replete but otherwise identical diet. After three hours of refeeding, the ILP2 content of IPCs dropped significantly, suggesting acute release of peptide into circulation (Fig. 1G). To confirm this, we measured circulating ILP2 levels [31]. When cholesterol-starved animals were transferred to food containing cholesterol, circulating hemolymph ILP2 levels increased slightly after one hour of cholesterol feeding and significantly after four hours (Fig. 1H). To assess the effect of cholesterol on peripheral insulin signaling, we measured the level of phosphorylation of Akt (pAkt), a readout of intracellular insulin-pathway activity. Cholesterol feeding for four hours increased whole-animal pAkt levels (Fig. 1I and 1J), indicating increased systemic insulin-signaling activity. Together these findings show that dietary cholesterol promotes insulin expression and increases the release of ILP2 and −5 into the hemolymph, thereby systemically promoting body growth.

### Cholesterol sensing promotes systemic growth mainly via glia and fat body

Next, we sought to identify the anatomical routes through which dietary cholesterol affects insulin. Although the IPCs are autonomously sensitive to some nutritional inputs, most nutrition-induced effects on these cells are mediated by sensory mechanisms in peripheral tissues. The fat body is a central nutrient-sensing hub [32], releasing multiple signals that govern the expression and release of ILPs [33]. Several neuronal cell types within the larval CNS have also been identified as modulators of the IPCs. Furthermore, the *Drosophila* nervous system, like that of mammals, is also surrounded by a selectively permeable layer of tightly apposed cells that form a BBB. In *Drosophila*, the BBB comprises nutrient-sensitive glial cells that relay amino-acid information to the IPCs to regulate ILP expression [11].

To assess whether any of these tissues or cells might mediate the effects of dietary cholesterol, we targeted them using tissue-specific GAL4 drivers: *Ilp2-GAL4* (the IPCs), *R57C10-GAL4* (pan-neuronal), *moody-GAL4* (BBB glia), and *Cg-GAL4* (fat body). To disrupt cholesterol trafficking and sensing in these cells, we used RNAi to knock down the expression of *Npc1a* or *Npc1b*, the *Drosophila* orthologues of *Npc1*. Npc1a is broadly expressed throughout the larva, whereas Npc1b is strongly expressed only in the gut. These paralogues are non-interchangeable and non-redundant [34], so we assessed the effects of knocking them down individually in each tissue. This manipulation should lead to high levels of intracellular cholesterol and increased Npc1-mediated cholesterol sensing [24]; thus, we aimed to identify tissues in which *Npc1a/b* RNAi brings about a body-size ***increase***, thereby phenocopying the response to dietary cholesterol supplementation.

Knockdown of either *Npc1a* or -*1b* in the IPCs did not significantly alter developmental timing, but reduced pupal size (Fig. 2A and 2A’), suggesting these cells do not drive cholesterol-induced growth. We also further excluded the involvement of ecdysone through manipulating the PTTH-producing neurons (PTTHn) [35]. Knockdown of *Npc1a* or -*1b* in these neurons using *Ptth-GAL4* did not significantly alter pupation timing or pupal size (Fig. S2A), consistent with cholesterol-induced growth’s being independent of ecdysone (Fig. 1D and Fig. S1). We next targeted the entire nervous system with pan-neuronal knockdown of *Npc1a* or *Npc1b* using *R57C10-GAL4* (*R57C10*>). Although knockdown of *Npc1b* led to accelerated pupariation, neuronal knockdown of *Npc* genes did not increase pupal size (Fig. 2B and 2B’). Thus, it seems unlikely that neuronal cholesterol sensing through Npc1 underlies the increased ILP production and release mediating the systemic growth-promoting effects of cholesterol.

**FIGURE 2.**
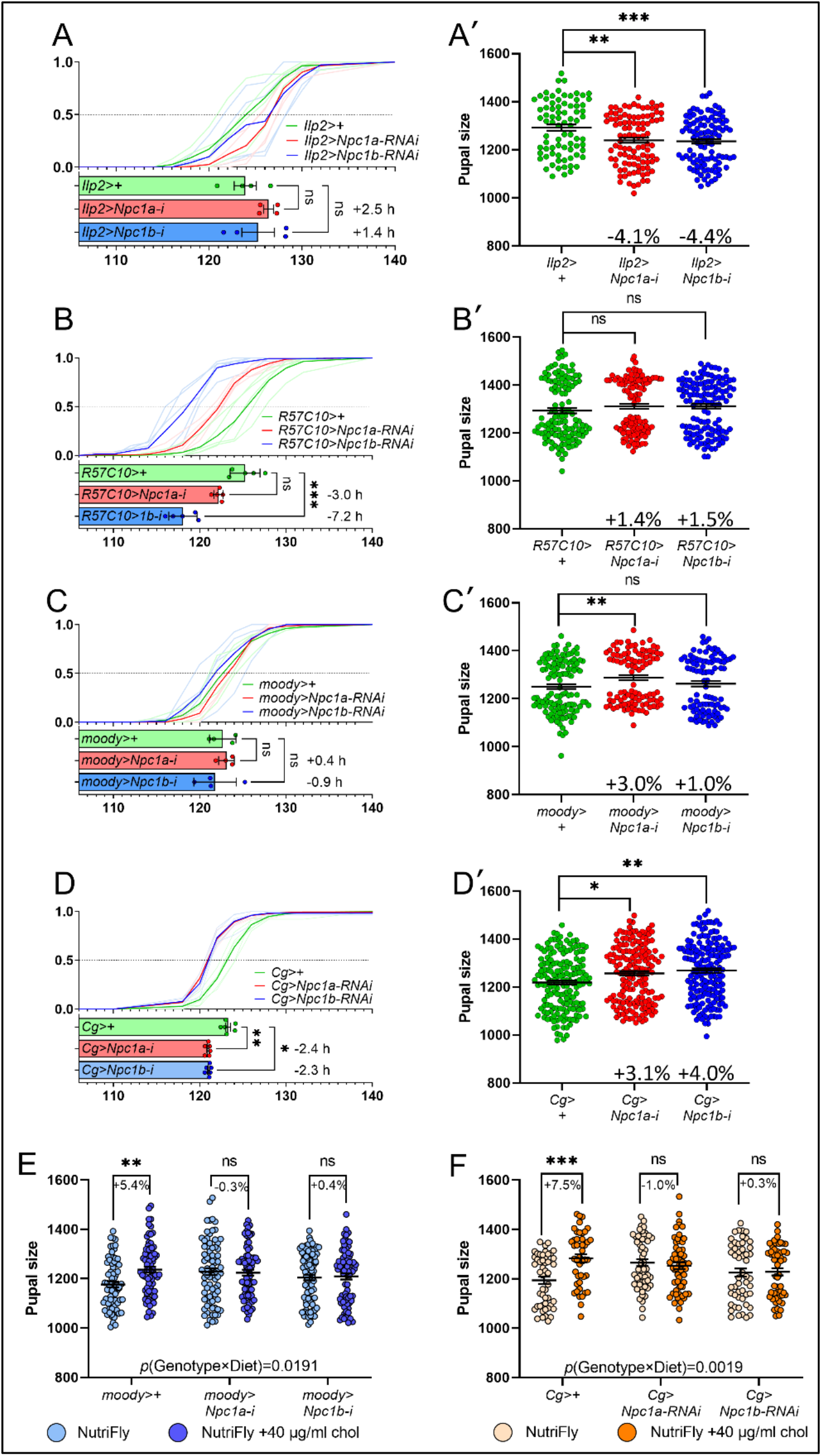
Cholesterol-induced growth promotion is mediated by the fat body and the glia of the blood-brain barrier. **A, B, C, D:** Top panel: pupariation over time on standard lab diet; individual trials are shown as fainter lines, and the average is shown in a darker line. Bottom panel: The time until 50% pupariation; each data point represents a vial of animals. **A’, B’, C’, D’:** Pupal sizes of animals from several vials of each genotype. In some cases, the sexual dimorphism in size is apparent as a bimodal distribution of points. **A**: Npc1 knockdown in the IPCs has no effect on developmental timing but reduces pupal size significantly. **B**: Knockdown of *Npc1* genes in the entire nervous system accelerates pupariation but has no significant effects on pupal size. **C**: Knockdown of *Npc1a* in the glia of the blood-brain barrier increases pupal size but has no effect on developmental timing; *Npc1b* knockdown has no effects on this diet. **D**: Knockdown of either Npc1a or Npc1b in the fat body accelerates pupariation and increases pupal size. **E** and **F**: *Npc1a/b* knockdown in the gli aof the BBB (**E**) or the fat body (**F**) is epistatic to dietary cholesterol. Leftmost pairs: Control animals in both tests fed on NutriFly medium supplemented with 40 μg/ml cholesterol (darker data points) are larger than animals fed unsupplemented NutriFly diet (lighter data points). Middle and right pair: knockdown of *Npc1a* or *Npc1b* in these tissues increases body size, and cholesterol supplementation has no additive effect, suggesting that cholesterol acts through a pathway containing Npc1a. Two-way ANOVA indicates strong genotype-by-diet interaction in both cases. Statistics: A-D’, Kruskal-Wallis ANOVA with Dunn’s multiple comparisons. E and F: Two-way ANOVA with Šídák’s multiple comparisons. Significance is noted as *, *p*<0.05; **, *p*<0.01; ***, *p*<0.001.

We next investigated whether BBB-glial and fat-body cholesterol levels might mediate the effects of cholesterol on insulin signaling and growth, since these tissues relay other types of nutritional information to the IPCs [9, 11]. Disrupting cholesterol trafficking in the BBB did not alter developmental timing, but *Npc1a* knockdown led to a significant increase in pupal size (Fig. 2C and 2C’). Altering cholesterol signaling in the fat body through knockdown of *Npc1a* or ™*1b* led to accelerated pupariation as well as increased pupal size (Fig. 2D and 2D’). Together these results indicate that increased cholesterol sensing in glia and fat body increases growth rate.

We next wished to determine the epistatic relationship between *Npc1a* knockdown and cholesterol availability to define whether this Npc1-mediated effect is cholesterol-specific (Fig. 2E and 2F). Cholesterol strongly increased the size of control pupae, whereas the increased size observed in animals with *Npc1*-gene knockdown in glia or fat body was not further enhanced by cholesterol supplementation (Fig. 2E and 2F). Similarly, increased dietary cholesterol accelerated the pupariation of control animals by several hours, but had little effect on the developmental timing of animals with BBB-glia or fat-body *Npc1a* gene knockdown (Fig. S2, B and C). Thus, Npc1a knockdown is epistatic to cholesterol level; if cholesterol and Npc1 knockdown were independent, we would expect an additive relationship, and thus these results suggest that the growth-accelerating effects of *Npc1a* knockdown in glia and the fat body are cholesterol-dependent. Taken together, our data suggest that the BBB glia and cells of the fat body sense cholesterol availability through a mechanism affected by Npc1 and regulate body growth accordingly.

### Cholesterol affects insulin signaling via glia and fat body relays

We next characterized the effects of cholesterol-trafficking/-sensing manipulations on insulin expression and release. Consistent with inability of IPC or PTTHn knockdown of *Npc1* genes to recapitulate the overgrowth phenotype, we found little evidence that these manipulations affect ILP or PTTH signaling. In animals with IPC knockdown of *Npc1* genes, IPC staining was unchanged, and the expression of *ILP*s and genes suppressed by insulin-pathway activity did not indicate increased insulin signaling (Fig. 3A and 3A’). This suggests cholesterol does not promote growth through Npc1-regulated sensing in the IPCs. Furthermore, PTTH staining also indicates that *Npc1a* knockdown in the PTTHn did not alter PTTH release (Fig. S2C). Knockdown of *Npc1a* or −*1b* in the entire nervous system with *R57C10*> led to reduced *4EBP* and *InR* transcript levels (Fig. 3B and 3B’), suggesting higher peripheral insulin signaling, consistent with the faster development seen in these animals. However, expression of *Ilp2* and −*3* was also significantly reduced, inconsistent with these effects, and the IPC levels of ILP2 and ILP5 staining were reduced as well. Thus, a minor influence of neuronal cholesterol signaling on larval growth cannot be excluded.

**FIGURE 3.**
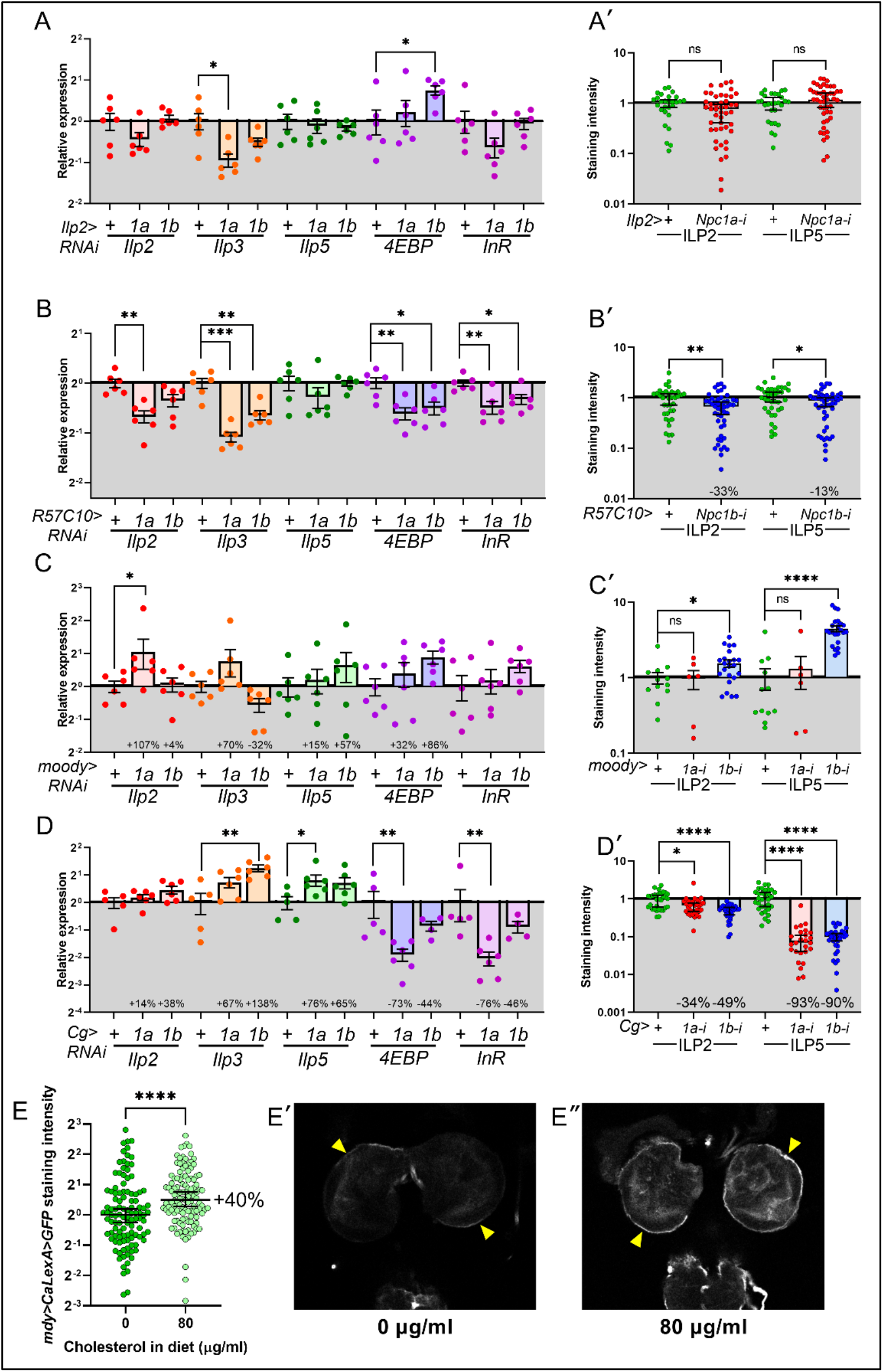
Manipulation of the cholesterol trafficking and signaling in the fat body and the BBB alter insulin expression and release. **A, B, C, D:** Gene expression profiles by qPCR of whole third-instar larvae. **A’, B’, C’, D’:** IPC stains of ILP2 and ILP5 as an indirect indicator of retention or release. **A/A’**: Knockdown of *Npc1a* in the IPCs has reduces *Ilp3* expression, and knockdown of *Npc1b* increases *4EBP* expression; no effect on IPC staining was observed. **B/B’**: *Npc1* knockdown in the IPCs appears to reduce *Ilp2/3* expression and ILP2/5 staining, but also reduces the expression of genes inhibited by insulin signaling (*4EBP* and *InR*), so the impact of these manipulations is unclear. **C/C’**: *Npc1a* knockdown in the BBB significantly increases *Ilp2* expression but does not alter IPC staining, suggesting increased ILP2 release. **D/D’**: *Npc1b* knockdown in the fat body significantly increases *Ilp3* expression, and *Npc1a* knockdown increases *Ilp5*; expression of *4EBP* and *InR* are significantly decreased by *Npc1a* knockdown, suggesting increased insulin signaling. Consistent with this, *Npc1* knockdown leads to strongly reduced ILP2 and ILP5 staining. **E:** Animals starved of cholesterol exhibit low levels of calcium-signaling activity in the cells of the BBB, as indicated by UAS-CaLexA> GFP staining; animals fed a diet containing 80 μg/ml cholesterol but otherwise identical exhibit increased BBB-glia activity. **E’ and Eʺ:** Illustrative images from the two treatments. BBB is marked with arrowheads. Statistics: A-D: Welch’s ANOVA with Dunnett’s T3 multiple comparisons for normally distributed data, Kruskal-Wallis test with Dunn’s multiple comparisons for non-normal data. A’-D’: Kruskal-Wallis test with Dunn’s multiple comparisons. E: Mann-Whitney test. Significance is noted as *, *p*<0.05; **, *p*<0.01; ***, *p*<0.001; ****, *p*<0.0001.

We then asked whether the increased size caused by *Npc1* gene knockdown in BBB glia and the fat body was associated with altered insulin signaling. Knockdown of *Npc1a* in the BBB significantly increased the expression of *Ilp2* and *Ilp3* (Fig. 3C and 3C’), consistent with previous reports suggesting that glia-mediated effects on ILPs are mainly transcriptional [11]. Next, we investigated whether cholesterol affected glial calcium signaling, which has previously been linked to mechanisms by which nutritional lipids modulate IPC activity [36]. Consistent with glial effects of cholesterol on systemic growth, we found that CaLexA, a reporter of intracellular calcium, in the BBB increased with cholesterol feeding (Fig. 3E). This suggests that cholesterol promotes glial cell activity that relays information to the IPCs to regulate their activity.

Increasing fat-body intracellular cholesterol by *Npc1a* knockdown led to a significant increase in *Ilp5* expression as well as strong reductions of ILP2 and ILP5 staining in the IPCs (Fig. 3D and 3D’), suggesting increased insulin release. Consistent with this and with the observed enhanced growth (Fig. 2D and 2D’), we also found strong reductions in the transcript levels of *4EBP* and *InR*, indicating increased systemic insulin signaling. We therefore examined whole-body expression levels of known fat-derived insulin-regulating factors and found that of these only *Ilp6* was altered by fat-body *Npc1a* knockdown (Fig. S3). *Ilp6* expression in the fat body itself is induced by starvation and ecdysone signaling [37, 38], and thus, the observed increase may not necessarily be involved in relaying cholesterol status from the fat body to the IPCs, but may rather be affected by downstream events.

### Enhanced cholesterol signaling in the PG alters the nutritional checkpoint that prevents precocious initiation of maturation

Nutrition influences ecdysone production through insulin and PTTH as well as TOR signaling in the PG itself [13, 30, 39, 40]. The TOR-mediated mechanism in the PG couples nutritional inputs to endocycling, which in turn is linked to CW [41, 42]. We therefore asked whether cholesterol levels in the PG affect these events. To observe the influence of cholesterol on endoreduplication, we used *GAL80^TS^; phm*> (*phm^ts^*>) to induce *Npc1a-RNAi* in the PG in the mid-second-instar stage (L2). This manipulation, which leads to strong intracellular cholesterol accumulation [29, 43], led to a massive increase in endoreduplication, reflected in increased nuclear size (Fig. 4A), suggesting that cholesterol sensing in this tissue is linked to this mechanism by which larvae estimate CW.

**FIGURE 4.**
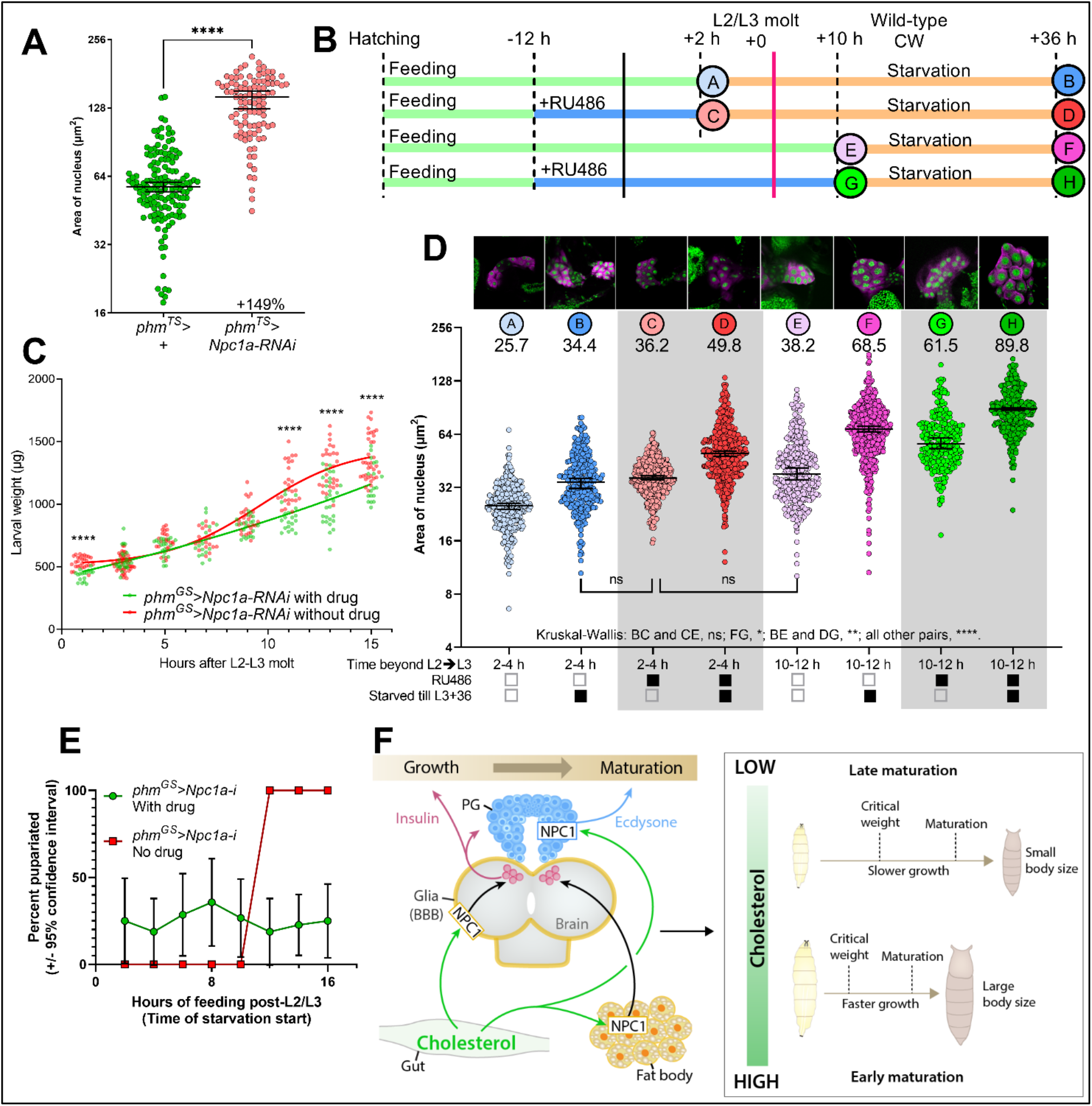
Increased cholesterol signaling in the PG promotes endoreduplication and alters the critical-weight checkpoint. **A:** Temperature-induced knockdown of *Npc1a* in the PG during third-instar life promotes endureduplication, as indicated by larger PG-cell nuclei. **B:** Schematic of the time course and the samples taken during the critical-weight experiment. **C:** Drug induction (green) of *phm-GeneSwitchGAL4*>*Npc1a-RNAi* reduces body growth at later times, compared to non-induced isogenic animals (red), suggesting that Npc1a knockdown increases ecdysone production. **D:** The nuclear size of PG cells increases over developmental time, and induction of Npc1a knockdown using RU486 promotes endureduplication as reflected in additional size increase. Note that PG nuclei of drug-induced animals examined at 2 hours after the L3 molt (sample C) are of the same size as the nuclei of non-induced animals allowed to feed for 10-12 hours (sample E). **E:** Consistent with their increased nuclear size, animals with *Npc1a* knockdown in the PG are able to pupariate immediately after the L3 molt (green circles), whereas non-induced control animals require 10-12 hours of feeding to attain CW, illustrated by their transition from 0% pupariation to 100% pupariation. **F:** Graphical model of findings. Statistics: A: Median with 95% confidence interval; Mann-Whitney test. C: Multiple unpaired t test with Welch correction, false discovery rate = 1%. D: Median with 95% confidence interval; Kruskal-Wallis test with Dunn’s multiple comparisons. E: Fraction of pupariated animals with margin of error, with each pupa representing a pupated/not-pupated value. Significance is noted as *, *p*<0.05; **, *p*<0.01; ***, *p*<0.001; ****, *p*<0.0001.

Since cholesterol sensing appears to drive endureduplication, and endoreduplication drives the irreversible activation of the pupariation-inducing neuroendocrine axis at CW [41, 42], we therefore investigated whether increasing PG cholesterol by knockdown of *Npc1a* would reduce CW – that is, if it might cause animals to overestimate their overall body size and therefore pupariate inappropriately. We used the drug-inducible *phantom-GeneSwitch-GAL4* (*phm^GS^*>) driver to induce *Npc1a-RNAi* in the PG in mid-L2 by treatment with dietary RU486. We measured PG-cell nuclear size in drug-treated animals versus non-treated controls after 2 or 10 hours of feeding after the L2-L3 molt, and after similar animals were starved until 36 hours post-molt (schematized in Fig. 4B). The body growth of animals with PG-specific *Npc1a* knockdown was attenuated in larvae beginning around 12-14 hours after the L2 to L3 transition (Figure 4C), consistent with an increase in ecdysone production, which inhibits systemic growth. *Npc1* knockdown in the PG led to a large increase in nuclear size at all time points tested (Figure 4D), indicative of increased endoreduplication. Indeed, PG nuclei were of similar sizes in newly molted L3 animals treated with RU486 (condition “C”) and in non-drug-treated controls at 10-12 hours (condition “E”), which had already attained CW as demonstrated by their 100% pupariation rate under starvation (Fig. 4E).

Control animals starved before 10 hours after the L2-L3 transition never pupariated, whereas all controls starved after this time pupariated, indicating that larvae attain CW after ~10 hours’ L3 feeding, at around 0.8 mg, in agreement with previous findings [44]. However, we found that a significant fraction of treated animals (with *Npc1a* knockdown in the PG) were able to pupariate when starved at early time points, even immediately following the L2-L3 molt (0-2 hours), indicating that they had passed CW, consistent with their increased endoreduplication. Treated animals as small as 0.5 mg pupariated without further feeding. Treated animals never attained a 100% pupariation rate, likely because *Npc1a* is important later in the L3 for the trafficking of cholesterol for the peak of ecdysone biosynthesis that triggers pupariation [29]. Nonetheless, our combined results show that *Npc1a* knockdown in the PG strongly increased endoreduplication and permitted inappropriate pupariation, indicating that it causes premature CW attainment. Together this shows that increasing PG cholesterol levels and sensing via *Npc1a* knockdown causes a strong reduction in the body size required to pass the nutritional CW checkpoint and thus to initiate nutrient-independent maturation.

## DISCUSSION

Nutrition is one of the most important influences on developmental growth and maturation. Malnutrition or disease can impair growth and delay puberty, whereas obese children enter puberty early [2]. Similarly, *Drosophila* larvae exposed to poor nutrition, tissue damage, or inflammation delay their development, while rich conditions promote rapid growth and maturation [1, 45–47]. These environmental factors are coupled to the appropriate production of steroids via internal checkpoints, one of which is a nutritional-dependent critical weight required to initiate the maturation process [8]. This suggests that signals reflecting nutritional state and body-fat storage play a key role in activating the neuroendocrine pathways that trigger puberty. Although studies suggest that the adipokine leptin may be involved [48], the mechanisms linking body fat to puberty are poorly defined, and the potential involvement of lifestyle associated with excessive accumulation of cholesterol, one of the most important lipids, has not been considered. In humans, white adipose tissue is the main site of cholesterol storage and can contain over half the body’s total cholesterol in obesity [49]. Our results show that dietary cholesterol intake promotes systemic body growth through insulin-dependent pathways and that animals raised on high dietary cholesterol initiate maturation earlier. We show that cholesterol is sensed through an Npc1-regulated mechanism in the fat body and the glial cells of the BBB, which relay information to the IPCs of the brain to promote insulin expression and release, thus coupling growth and maturation with cholesterol status.

Insect CW evolved to ensure that maturation will not occur unless the animal has accumulated adequate nutrient stores to survive the non-feeding metamorphosis period and has completed sufficient growth to produce an adult of proper size and thus of maximal fitness. Likewise, the link between body fat and maturation in humans probably ensures adequate stores of fat before maturation onset to support pregnancy and reproductive success [17]. In *Drosophila*, insulin signaling plays a critical role in coordinating steroidogenesis with nutritional conditions. Insulin acts upon the PG and induces a small ecdysone peak early in L3 that is correlated with CW attainment [44, 50]. In combination with nutrition-related signaling mediated via insulin, nutrient availability is also assessed directly in the PG and is coupled to irreversible endoreduplication that permits ecdysone production at CW [41, 42]. Our findings show that accumulation of cholesterol in the PG, induced by loss of *Npc1a*, drives a remarkable increase in endoreduplication and precocious attainment of the CW.

Although neurodegeneration is the hallmark of NPC disease, including in a *Drosophila* model [51], alterations in glial, adipose, hepatic, and endocrinological systems are also components of NPC syndrome. In humans Npc1 itself is strongly expressed in glia [52] and in adipose tissues, especially in obese individuals [53], and variants of Npc1 are associated with obesity, type-2 diabetes, and hepatic lipid dysfunction [54–58]. Our findings link glial and adipose-tissue cholesterol sensing through Npc1 to systemic growth and metabolic control through effects on insulin signaling. We also find that intracellular cholesterol accumulation driven by *Npc* loss leads to increases in DNA replication and cell growth, which provides mechanistic insight for understanding the emerging link between cholesterol and cancers [59].

As the coupling of nutrition with growth and maturation is ancient and highly conserved, our findings provide a foundation for understanding how cholesterol is coupled to developmental growth and maturation initiation in humans. Our finding links a high concentration of this particular lipid in adipose tissues to neuroendocrine initiation of maturation, which may explain the critical link between obesity (body fat) and early puberty.

## MATERIALS AND METHODS

### Fly media and husbandry

For general stock-keeping, flies were kept on medium containing 6% sucrose, 0.8% agar, 3.4% yeast, 8.2% cornmeal, 0.16% Tegosept antifungal, and 0.48% propionic acid preservative. The synthetic diet used in the experiments of Fig. 1A and 1B was that described by Reis *et al*. [25]. This medium appears to be nutritionally poor and is very liquid, so the remaining experiments involving completely synthetic diet used the Reis recipe with the following adjustments: casein was lipid-depleted three times for at least 4 hours each by stirring with 5 volumes of chloroform to remove any lingering sterols, followed by filtration and drying, and its concentration was doubled. Agar was similarly lipid-depleted and added to a 1% final concentration. Sucrose was increased 2.7-fold. Choline chloride was doubled. In either case, cholesterol was added as 100x solutions in 100% ethanol (100 μl into each 10-ml vial; pure ethanol was added for cholesterol-free diet). Other experiments made use of pre-mixed NutriFly medium (Genesee Scientific, “Bloomington formula”), which has a relatively low yeast/cholesterol content, either as-is or supplemented with cholesterol: vials containing 10 ml prepared NutriFly medium were re-liquefied by microwaving, 50 or 100 μl of 100% ethanol or ethanol containing 8 mg/ml cholesterol was added to reach 0, 40, or 80 μg/ml final cholesterol concentration, and the medium was thoroughly mixed. Flies were kept at 25 °C and 60% humidity with a 12-hour/12-hour daily light cycle unless otherwise noted.

### Fly stocks

Stocks obtained from the University of Indiana Bloomington *Drosophila* Stock Center (BDSC) include *Cg-GAL4* (#7011), *moody-GAL4* [60] (#90883), *R57C10-GAL4* [61] (an *nSyb-GAL4* variant from Janelia Research Campus, #39171), and *UAS-Calcium-LexA*>*GFP* [62] (“CaLexA”, #66542). The Vienna *Drosophila* Resource Center (VDRC) provided lines including *UAS-Npc1a-RNAi* (#105405) and *UAS-Npc1b-RNAi* (#108054). M. B. O’Connor (University of Minnesota) kindly provided *phm-GAL4* [63], *phm-GeneSwitch-GAL4* [64], and *Ptth-GAL4* [35]. *Ilp2-GAL4, UAS-GFP* carries the *Ilp2-GAL4* construct described by Rulifson *et al.* [27], available separately as BDSC stock #37516; this and *Ptth-GAL4, UAS-Dicer-2* were gifts from Pierre Léopold (Institut Curie). *UAS-DILP2HF* [31] was a kind gift of S. Park and S. Kim (Stanford).

### Timed egg-lays

Virgin females of the driver stocks were mated with males of the RNAi lines in egg-lay chambers with apple-juice/agar egg-lay plates supplemented with yeast paste. Eggs were laid between ZT 1 and 5 (*i.e.*, one hour to five hours after incubator lights-on) at room temperature. For genotypes including *tub-GAL80^TS^*, eggs were allowed to develop on the plates for 48 hours at 18°, and first-instar larvae were collected and transferred to food vials (~30 per vial); transferred larvae were incubated a further 48 hours at 18° to prevent GAL40 activity during early larval development, prior to incubation at 29°. Larvae of other genotypes were collected after 24 hours’ development at 25 °C, transferred to vials (~30 per vial), and incubated at 25°.

### Developmental timing (pupariation timing)

Timed egg-lays of the appropriate genotypes were made, and animals that had begun to pupate were counted every 2-8 hours. The salient feature for the purpose of this measurement was the darkening of the pupal cuticle.

### Pupal-size measurement

After all animals in a vial had pupariated, animals were rinsed from the vial with deionized water and arranged on a glass microscope slide. Slides were imaged using a USB camera (Point Grey Grasshopper3, FLIR Systems, Inc.) and software (Point Grey FlyCapture, FLIR Systems, Inc.), and pupal sizes were determined from the images using a script[15] in the MATLAB environment (The MathWorks, Inc.).

### Quantitative real-time PCR (qPCR)

Six replicate RNA preparations for each genotype or condition were prepared using the NucleoSpin RNA Plus Mini kit (Macherey-Nagel, #740984.50) kit; each sample contained 3-5 whole late feeding third-instar larvae or 5 larval brains. Animals or tissues were placed in 2-mL Eppendorf tubes containing lysis buffer + 1% beta-mercaptoethanol, and samples were lysed using a bead mill (Qiagen). RNA was purified according to the kit instructions, and cDNA was prepared using the High-Capacity cDNA Synthesis Kit (ThermoFisher, #4368814). QPCR reactions were prepared using RealQ Plus 2x Master Mix Green without ROX (Ampliqon, #A323402) and the gene-specific primers given in the STAR methods, and runs were performed using a QuantStudio 5 machine (Applied Biosystems).

### Western blotting

Three feeding late-third-instar larvae for each sample were lysed in 3x SDS loading buffer (Bio-Rad) + 5% beta-mercaptoethanol using a bead mill (Qiagen). Samples were heated at 95° for 5 minutes to denature proteins, and insoluble material was pelleted by centrifugation at top speed for 1 minute. Proteins were separated on a precast 4%−20% gradient polyacrylamide gel (Bio-Rad)and transferred to PVDF membrane (Millipore) using a Trans-Blot Turbo (Bio-Rad) dry-transfer apparatus. The blot was blocked in Odyssey blocking buffer (LI-COR) for two hours at 4° with gentle agitation. Phospho-Akt and histone H3 were detected by incubation with rabbit anti-phospho-Akt (Cell Signaling Technologies, #4054, diluted 1:1000) and rabbit anti-histone-H3 (Abcam #1791, diluted 1:1000) in Odyssey blocking buffer + 0.2% Tween 20 (Sigma). (Akt and H3 are of very different sizes, allowing them to be detected together.) The blot was followed by 3 rinses in PBS+0.1% Tween 20 and secondary staining with IRDye 680RD-labeled goat anti-mouse and IRDye 800CW goat anti-rabbit (LI-COR, #925-68070 and #925-32210, 1:10,000 dilution in Odyssey blocking buffer + 0.2% Tween 20). Bands were visualized using an Odyssey Fc gel reader (LI-COR). The blot was stripped using NewBlot IR Stripping Buffer (LI-COR, #928-40028), and rinsed 3x with PBS + 0.1% Tween 20. Total Akt was detected on the stripped blot with rabbit anti-pan-Akt (Cell Signaling Technologies, #4691).

### Anti-DILP5 preparation

Antibodies were raised in rats against the peptide CPNGFNSMFAK, an epitope within the B chain of DILP5, and affinity-purified against the peptide by Genosys, Inc.

### Immunostaining, microscopy, and quantification

Tissues were dissected in cold PBS and fixed in fresh 4% paraformaldehyde in PBS at room temperature for one hour. For antibody staining, samples were quickly rinsed in PBST [PBS + 0.1% Triton X-100 (Sigma)], washed three times for fifteen minutes in PBST, and blocked for at least an hour at 4° in PBST + 3% normal goat serum (Sigma). Tissues were incubated overnight at 4° with gentle agitation with primary antibodies diluted in PBST and washed three times with PBST. Samples were incubated with secondary antibodies diluted in PBST at 4° overnight with gentle agitation and washed three times with PBST. Tissues were mounted on glass slides or glass-bottomed-dishes (MatTek Life Sciences, #P35G-1.5-10-C) coated with polylysine (Sigma, P8920-100ML) in ProLong Glass anti-fade mountant containing NucBlue DNA stain (ThermoFisher #P36985). Mounted tissues were imaged using a Zeiss LSM 900 confocal microscope using a 20x, NA 0.8 air-immersion objective. For quantification of staining, Z-stacks were collapsed in FIJI (NIH) using the “sum” method, IPC clusters were manually segmented, and the “raw integrated density” measurement was recorded. The region of interest was moved to an adjacent unstained area of tissue, and the “raw integrated density” of this background was measured and subtracted from the first measurement to give the net signal for each cell cluster. For the BBB CaLexA assay, linear regions of interest were drawn through the glial layer, perpendicular to the brain surface, in the plane of maximum brain size (in which the glial layer is most perpendicular to the image plane), using the line tool in FIJI; the intensity of CaLexA was measured along these lines, and the peak intensity along each line was recorded.

Primary antibodies included rabbit anti-DILP2 [65], diluted 1:1000; rabbit anti-DILP3 [66], a generous gift of Jan Veenstra (University of Bordeaux), diluted 1:500; rat anti-DILP5 (this work), diluted 1:500; mouse anti-GFP (clone 3E6, ThermoFisher #A-11120, RRID AB_221568), diluted 1:500; and guinea-pig anti-PTTH serum [67] (1:400), a kind gift of Pierre Léopold (Institut Curie).

Secondary antibodies were all raised in goats and were cross-adsorbed by the manufacturer to reduce off-target binding; they were obtained from ThermoFisher and diluted 1:500. These included anti-rabbit, Alexa Fluor 488 conjugate (#A32731, RRID AB_2633280), Alexa Fluor 555 conjugate (#A32732, RRID AB_2633281), and Alexa Fluor 647 conjugate (#A32733, RRID AB_2633282); anti-rat, Alexa Fluor 555 conjugate (#A48263, no RRID); anti-guinea pig, Alexa Fluor 647 conjugate (#A21450, RRID AB_2735091); and anti-mouse, Alexa Fluor 488 conjugate (#A32723, RRID AB_2633275).

### DILP2HF ELISA

*Ilp2>DILP2HF* [31] animals were reared on normal food until 80 hours after egg laying. At this time, they were transferred to cholesterol-free synthetic medium overnight (16 hours); after this sterol starvation, animals were transferred onto cholesterol-replete synthetic medium, and hemolymph was sampled at 0, 1, and 4 hours. Five-microliter batches of hemolymph were heat-treated to prevent coagulation and oxidation, and treated material was stored at −80 °C until use. “F8 MaxiSorp Nunc-Immuno modules” (Thermo Scientific #468667) were coated with anti-FLAG by incubating them overnight with 5 μg/mL mouse anti-FLAG (Sigma #F1804) in 200-mM NaHCO_3_ buffer (pH 9.4) at 4 °C. Wells were washed twice with PBS+0.1% Triton X-100 (PBST), blocked with PBST+4% non-fat dry milk for 2 h at room temperature, and washed three more times in PBST. Anti-HA::peroxidase (Roche #12013819001) was diluted to 25 ng/ml in PBST + 1% non-fat dry milk. Five microliters of hemolymph was added to 50 μL of this solution, and the mixture was incubated in the coated wells overnight at 4 °C. The wells were emptied and washed six times with PBST. The color reaction was started by adding 100 μL of One-step Ultra TMB ELISA substrate (Thermo Scientific #34028) was added to each well (100 μL); after sufficient color development (~1 minute), the reaction was stopped with the addition of 100 μL 2-M sulfuric acid. Absorbance at 450 nm was measured using a PerkinElmer EnSight multimode plate reader.

### Critical-weight determination

Embryos of *phm-GeneSwitchGAL4*>*Npc1a-RNAi* were collected on NutriFly egg-lay plates in a series of two-hour egg lays. At 52-54 hours AEL (mid-L2), the medium from each time point was cut in half, and the two halves were joined to half-plates of fresh NF medium. The rejoined halves, roughly 15 ml in total, were painted with 500 μl of a yeast suspension (10% w/v dried *S. cerevisiae* in water) containing either RU486 at 100 μg/ml (from a 50-mg/ml ethanol stock) or the equivalent volume of ethanol alone; the final concentration of drug in dosed plates, assuming even diffusion, was roughly 3 μg/ml. Animals were incubated at 25 °C for a further 12 hours, after which freshly molted L3 animals (determined based on anterior spiracle morphology) were harvested by flotation on 20% sucrose and manual sorting every two hours and transferred into vials containing NutriFly or NutriFly containing 3 μg/ml RU486. At designated time points, animals were re-harvested from these vials by flotation in 20% sucrose. Batches of animals were prepared for PG staining or for critical-weight assays. **For PG staining,** larvae were grossly dissected in PBS and fixed in fresh 4% paraformaldehyde in water at room temperature for precisely one hour with rotation. Tissue was rinsed with PBS+0.1% Triton X-100 (PBST) three times for 15 minutes each and blocked with PBST+3% normal goat serum at 4° overnight. Tissue was stained with rabbit anti-Phantom [63] (1:400), a kind gift of Michael O’Connor, at 4° overnight with gentle agitation, followed by three washes with PBST. Tissue was incubated at 4° overnight with goat anti-rabbit Alexa Fluor 555 (ThermoFisher #A32732, RRID AB_2633281, 1:500; Alexa Fluor 488-conjugated phalloidin (ThermoFisher #12379, 1:100), and DAPI (ThermoFisher #D1306, 1:500). Tissue was rinsed three times with PBST, equilibrated in 50% glycerol/water, and mounted on poly-lysine-coated MatTek imaging dishes in 50% glycerol/water. This liquid was withdrawn and replaced with ProLong Glass solidifying anti-fade mountant (ThermoFisher). After the mountant solidified, PGs were imaged as above. Each data point represents the cross-sectional area of a given nucleus at its greatest diameter, determined in FIJI: for each nucleus, the focal section demonstrating the greatest size was found; the nucleus was manually outlined, and its area was measured. **For critical-weight determination,** reharvested animals were individually weighed using a Sartorius SE2 ultramicro balance and transferred to starvation medium (0.6% agarose in water). The number of pupariated animals was recorded periodically. Some animals were also harvested from starvation medium at 36 hours after the L2/L3 molt and prepared for PG imaging.

## Supporting information

Supplemental table 1: qPCR oligo sequences

## Data availability and statistics

Data are available upon reasonable request. Statistics were computed and graphs were prepared in the Prism software package (GraphPad). Means are presented with the standard error. Data sets were assessed for normality prior to computation of statistics. Statistics are discussed in each legend. *P* values are represented in all figures as follows: **ns**, *p*>0.05; *, *p*≤0.05; **, *p*≤0.01; ***, *p*≤0.001; ****, *p*≤0.0001.

## Acknowledgements

We warmly thank our colleagues who have contributed materials, the use of equipment, and expertise to this effort: Jan Veenstra, Takashi Koyama, Pierre Léopold, Michael O’Connor, and the Bloomington and Vienna *Drosophila* stock centers. This work was supported by grant #2019-772 from the Danish Research Council, Natural Sciences, to KR.

## Author contributions

MJT and KR designed the study and experiments. MJT, KR, LHP, ML, and AM performed experiments and analyzed data. MJT and KR wrote the manuscript. KR obtained funding.

---

KEY RESOURCES TABLE.

KEY RESOURCES TABLE
| REAGENT or RESOURCE | SOURCE | IDENTIFIER |
| --- | --- | --- |
| **Antibodies** |  |  |
| Anti-Akt, phospho-Ser505, rabbit | Cell Signaling Technologies #4054 | (discontinued) |
| Anti-Akt, total, rabbit monoclonal C67E7 | Cell Signaling Technologies #4691 | No RRID |
| Anti-DILP2, rabbit [65] | E. Hafen (ETH Zurich)<br>M. Pankratz (Bonn) | No RRID |
| Anti-DILP3, rabbit [66] | J. Veenstra (Bordeaux) | No RRID |
| Anti-DILP5, rat | This paper | No RRID |
| Anti-FLAG, mouse clone M2 | Sigma #F1804 | No RRID |
| Anti-GFP, mouse clone 3E6 | ThermoFisher #A11120 | RRID AB_221568 |
| Anti-guinea pig IgG, Alexa Fluor 647 conjugate | ThermoFisher #A21450 | RRID AB_2735091 |
| Anti-HA, rat clone 3F10, peroxidase conjugate | Roche #12013819001 | No RRID |
| Anti-histone H3, rabbit | Abcam #1791 | No RRID |
| Anti-mouse IgG, Alexa Fluor 488 conjugate | ThermoFisher #A32723 | RRID AB_2633275 |
| Anti-mouse IgG, IRDye 680RD conjugate | LI-COR #925-68070 | RRID AB_2651128 |
| Anti-PTTH, guinea pig [67] | P. Léopold (Institut Curie) | No RRID |
| Anti-rabbit IgG, Alexa Fluor 488 conjugate | ThermoFisher #A32731 | RRID AB_2633280 |
| Anti-rabbit IgG, Alexa Fluor 555 conjugate | ThermoFisher #A32732 | RRID AB_2633281 |
| Anti-rabbit IgG, Alexa Fluor 647 conjugate | ThermoFisher #A32733 | RRID AB_2633282 |
| Anti-rabbit IgG, IRDye800CW conjugate | LI-COR #925-32210 | RRID AB_2687825 |
| Anti-rat, Alexa Fluor 555 conjugate | ThermoFisher #A48263 | No RRID |
| **Chemicals, peptides, and recombinant proteins** |  |  |
| Biotin | Sigma, #B4501 | CAS 58-85-5 |
| Casein | Sigma, #C5679 | CAS 9000-71-9 |
| Cholesterol | Sigma, #C3045 | CAS 57-88-5 |
| Choline chloride | Sigma, #C1879 | CAS 67-48-1 |
| Folic acid | Sigma, #F8758 | CAS 59-30-3 |
| Inosine | Sigma, #I4125 | CAS 58-63-9 |
| Nicotinic acid | Sigma, #N0761 | CAS 59-67-6 |
| D-Pantothenic acid, hemicalcium salt | Sigma, #P5155 | CAS 137-08-6 |
| Pyridoxine | Sigma, #P6280 | CAS 58-56-0 |
| Riboflavin | Sigma, #R4500 | CAS 83-88-5 |
| Thiamine | Sigma, #T1270 | CAS 67-03-8 |
| Uridine | Sigma, #U3003 | CAS 58-96-8 |
| **Commercial kits** |  |  |
| NucleoSpin RNA Plus Mini kit | Macherey-Nagel | #740984.50 |
| High-Capacity cDNA Synthesis kit | ThermoFisher | #4368814 |
| RealQ Plus 2x Master Mix Green without ROX | Ampliqon | #A323402 |
| **Organisms (*Drosophila* stocks)** |  |  |
| *Cg-GAL4* | BDSC #7011 |  |
| *Ilp2-LexA::VP16* [68] | N. Cordero, U. Nice |  |
| *Ilp2-GAL4, UAS-GFP* [27] | P. Léopold, Institut Curie |  |
| *LexAop-Kir2.1* [69] | L. Vasconcelos, Gulbenkian Inst. |  |
| *phm-GAL4* [63] | M. B. O'Connor, U. Minnesota |  |
| *Ptth-GAL4, UAS-Dicer-2* [35] | P. Léopold, Institut Curie |  |
| *R57C10-GAL4* [61] | BDSC #39171 |  |
| *SPG-GAL4* [60] | BDSC #90883 |  |
| *UAS-Calcium-LexA>GFP* [62] | BDSC #66542 |  |
| *UAS-Npc1a-RNAi* | VDRC #105405 |  |
| *UAS-Npc1b-RNAi* | VDRC #108054 |  |
| *w<sup>1118</sup>* | VDRC #60000 |  |
| Oligonucleotides |  |  |
| Please see Table S1 for qPCR oligo sequences |  |  |
| Software and algorithms |  |  |
| Pupal-size script [15] |  |  |

**Figure S1:**
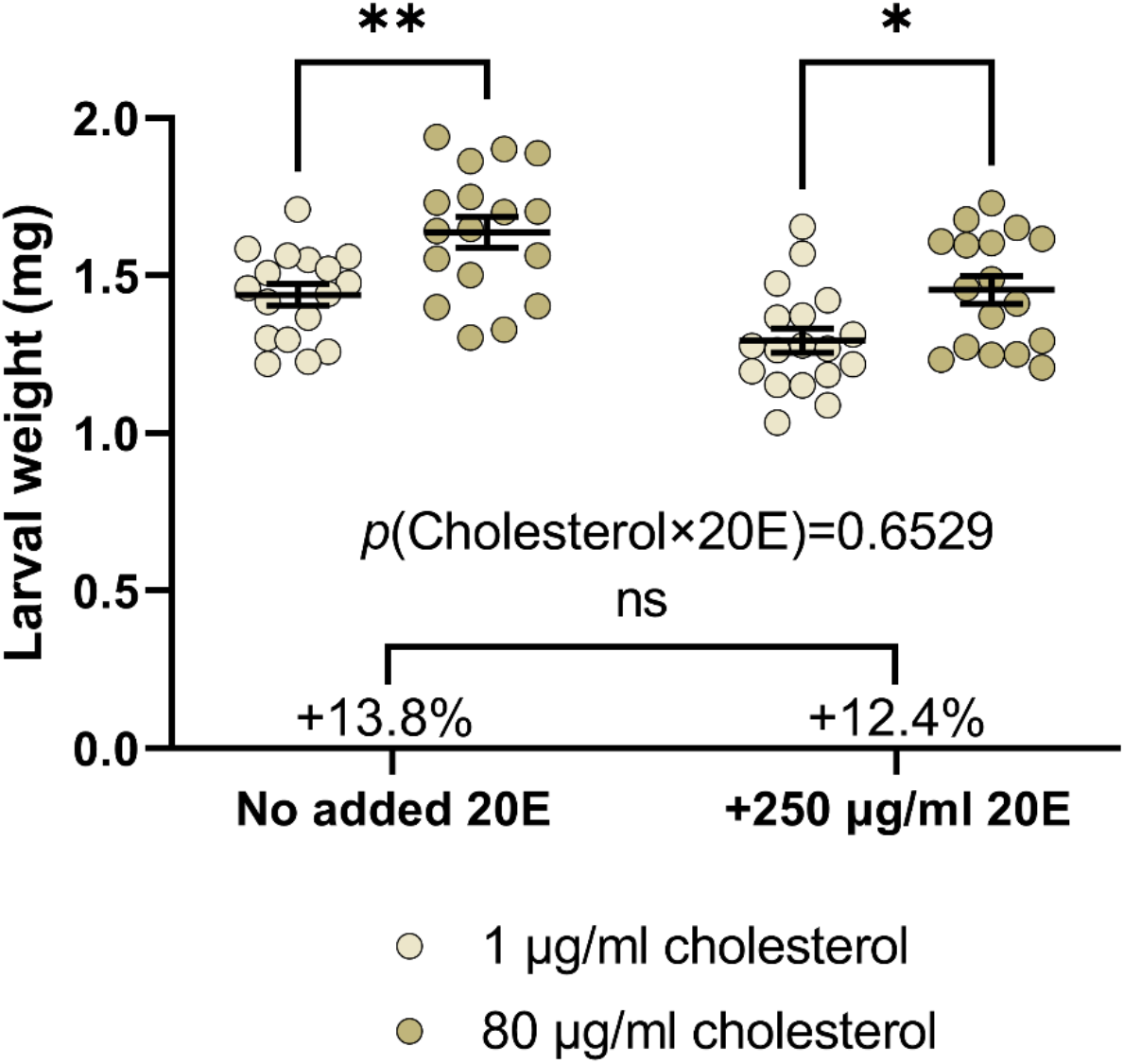
No interaction was observed between ecdysone and dietary cholesterol. Related to Figure 1D. Wild-type animals were fed on normal food, then transferred to synthetic food containing either 1 or 80 μg/ml cholesterol, with or without 250 μg/ml 20E. Higher cholesterol (darker points) promoted growth to a similar extent whether or not 20E was directly added. Two-way ANOVA with Šídák’s multiple comparisons; *p*=0.65 for cholesterol-by-ecdysone interaction. Significance is noted as *, p<0.05; **, p<0.01.

**Figure S2.**
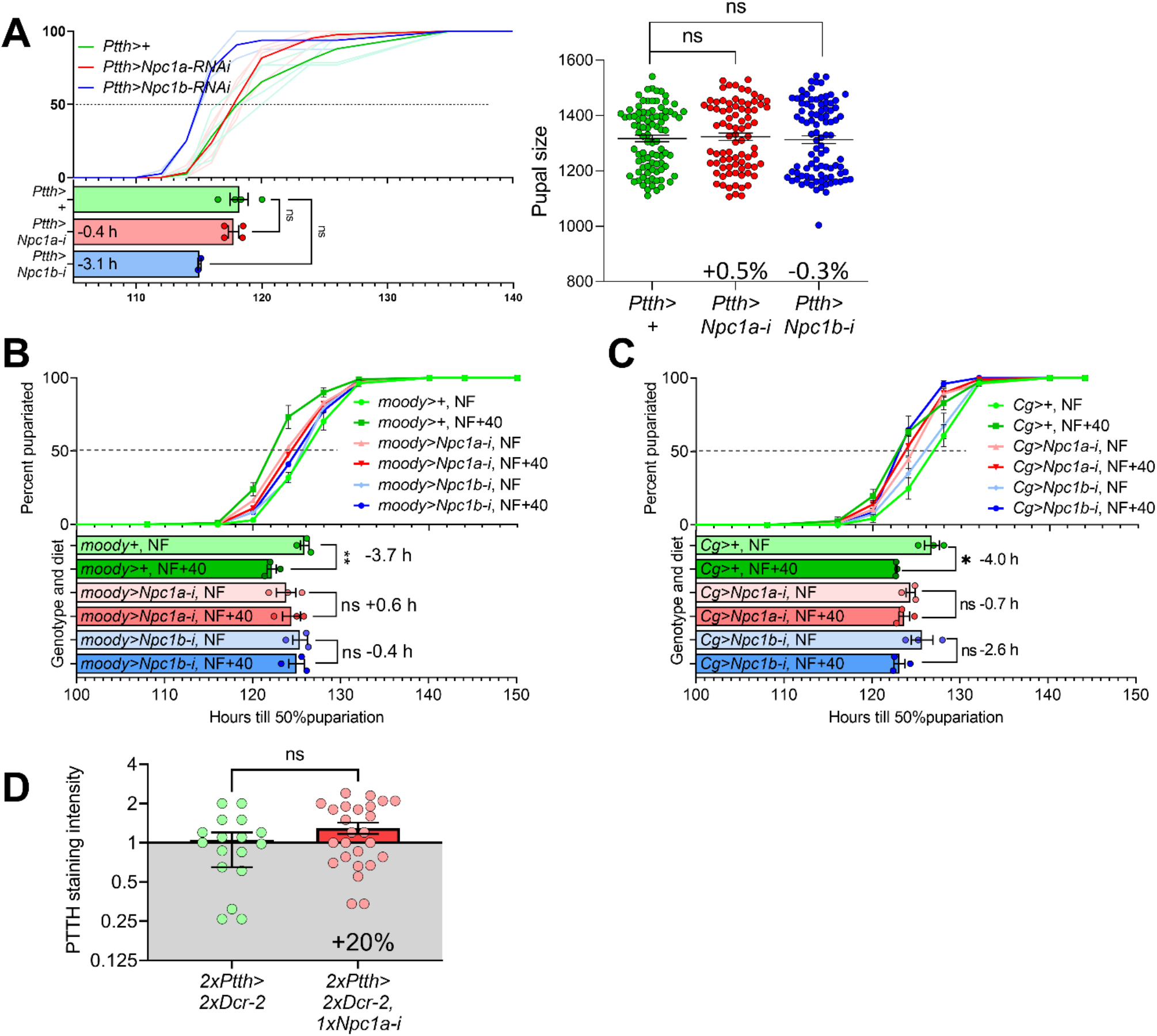
**A:** Supplement to Figure 2. *Npc1* RNAi in the PTTH-producing cells has no significant effect on developmental timing (left), and neither *Npc1* orthologue affects pupal size (right). **B and C:** Related to Fig. 2, E and F. *Npc1a* RNAi in the BBB glia **(B)** or the cells of the fat body **(C)** is epistatic to dietary cholesterol with regard to developmental acceleration. In control animals (*moody*>+ or *Cg*>+, green, top two bars), added cholesterol accelerated pupariation by ~4 hours; when *Npc1a* expression was knocked down (*moody*>*Npc1a-i* or *Cg>Npc1a-i*, red bars, middle two bars), cholesterol supplementation did not affect developmental timing, suggesting that cholesterol acts through an Npc1a-mediated process. **D:** Related to Fig. 1D, Fig. S1, and Fig. S2A. Knockdown of *Npc1a* in the PTTH-producing cells did not significantly alter PTTH staining intensity in these cells, consistent with the other results that indicate no role for PTTH or ecdysone in cholesterol-induced acceleration of growth and development. Statistics: A, as in Figure 2; B and C: unpaired t-test with Welch’s correction between NF and NF+40 for each genotype. D: Kolmogorov-Smirnov test. Significance is noted as *, p<0.05; **, p<0.01.

**Figure S3:**
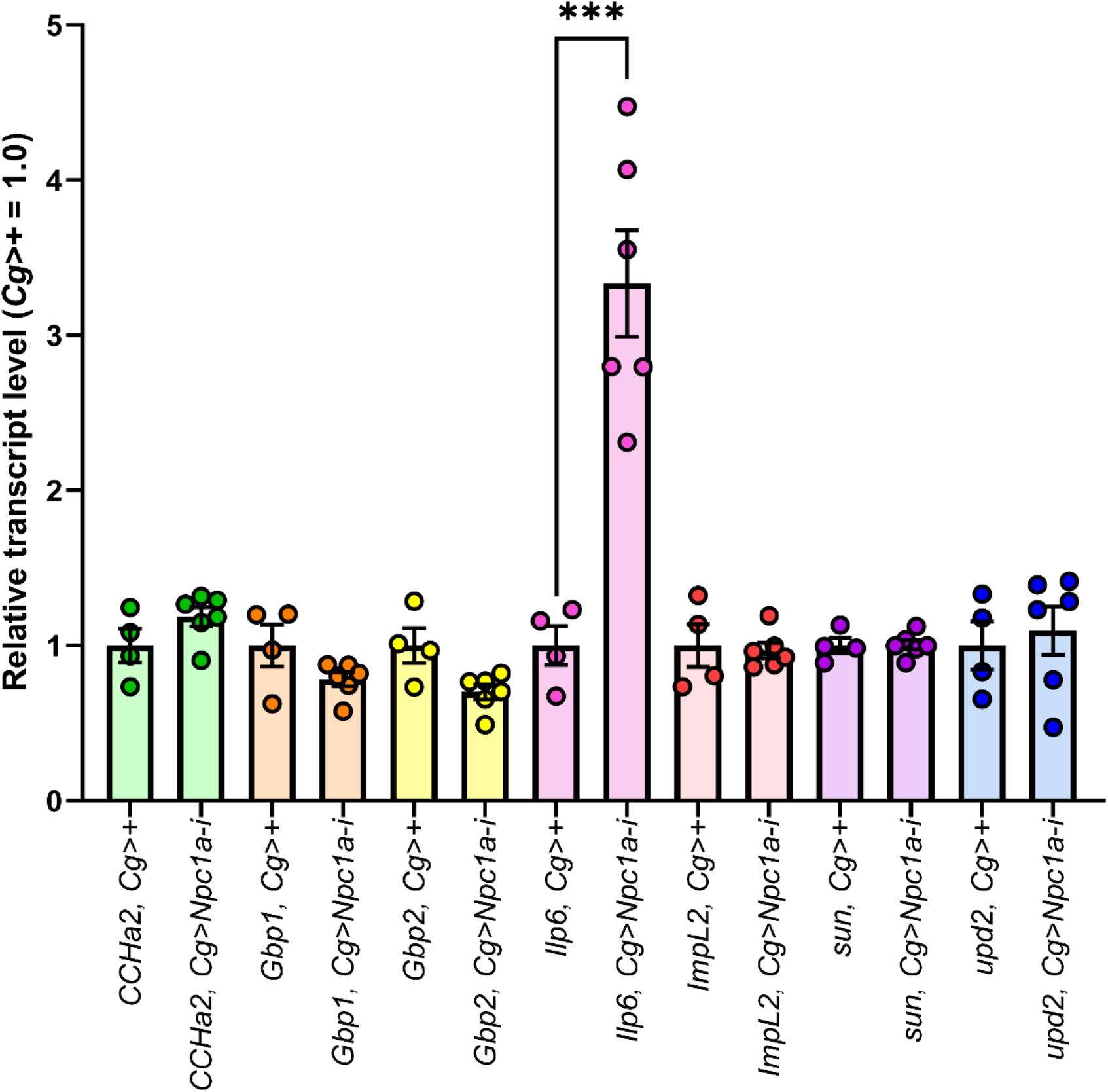
Npc1a RNAi in the fat body altered the transcript levels of *Ilp6* but not other tested fat-body-derived signals in whole-animal qPCR. Unpaired t tests with Welch’s correction for each target gene. Significance is noted as ***, *p*<0.001.

## REFERENCES

1. Texada, M.J., Koyama, T., and Rewitz, K. (2020). Regulation of Body Size and Growth Control. Genetics 216, 269–313.

2. Ahmed, M.L., Ong, K.K., and Dunger, D.B. (2009). Childhood obesity and the timing of puberty. Trends Endocrinol Metab 20, 237–242.

3. Navarro, V.M., Castellano, J.M., Garcia-Galiano, D., and Tena-Sempere, M. (2007). Neuroendocrine factors in the initiation of puberty: the emergent role of kisspeptin. Rev Endocr Metab Disord 8, 11–20.

4. Deveci, D., Martin, F.A., Leopold, P., and Romero, N.M. (2019). AstA Signaling Functions as an Evolutionary Conserved Mechanism Timing Juvenile to Adult Transition. Curr Biol 29, 813–822 e814.

5. Imura, E., Shimada-Niwa, Y., Nishimura, T., Huckesfeld, S., Schlegel, P., Ohhara, Y., Kondo, S., Tanimoto, H., Cardona, A., Pankratz, M.J., et al. (2020). The Corazonin-PTTH Neuronal Axis Controls Systemic Body Growth by Regulating Basal Ecdysteroid Biosynthesis in Drosophila melanogaster. Curr Biol 30, 2156–2165 e2155.

6. Christensen, C.F., Koyama, T., Nagy, S., Danielsen, E.T., Texada, M.J., Halberg, K.A., and Rewitz, K. (2020). Ecdysone-dependent feedback regulation of prothoracicotropic hormone controls the timing of developmental maturation. Development 147.

7. Moeller, M.E., Danielsen, E.T., Herder, R., O’Connor, M.B., and Rewitz, K.F. (2013). Dynamic feedback circuits function as a switch for shaping a maturation-inducing steroid pulse in Drosophila. Development 140, 4730–4739.

8. Malita, A., and Rewitz, K. (2021). Interorgan communication in the control of metamorphosis. Curr Opin Insect Sci 43, 54–62.

9. Colombani, J., Raisin, S., Pantalacci, S., Radimerski, T., Montagne, J., and Leopold, P. (2003). A nutrient sensor mechanism controls Drosophila growth. Cell 114, 739–749.

10. Texada, M., Koyama, T., and Rewitz, K. (2020). Regulation of Body Size and Growth Control. Genetics In press.

11. Okamoto, N., and Nishimura, T. (2015). Signaling from Glia and Cholinergic Neurons Controls Nutrient-Dependent Production of an Insulin-like Peptide for Drosophila Body Growth. Dev Cell 35, 295–310.

12. Caldwell, P.E., Walkiewicz, M., and Stern, M. (2005). Ras activity in the Drosophila prothoracic gland regulates body size and developmental rate via ecdysone release. Curr Biol 15, 1785–1795.

13. Colombani, J., Bianchini, L., Layalle, S., Pondeville, E., Dauphin-Villemant, C., Antoniewski, C., Carre, C., Noselli, S., and Leopold, P. (2005). Antagonistic actions of ecdysone and insulins determine final size in Drosophila. Science 310, 667–670.

14. Boulan, L., Martin, D., and Milan, M. (2013). bantam miRNA promotes systemic growth by connecting insulin signaling and ecdysone production. Curr Biol 23, 473–478.

15. Moeller, M.E., Nagy, S., Gerlach, S.U., Soegaard, K.C., Danielsen, E.T., Texada, M.J., and Rewitz, K.F. (2017). Warts Signaling Controls Organ and Body Growth through Regulation of Ecdysone. Curr Biol 27, 1652–1659 e1654.

16. Texada, M.J., Malita, A., and Rewitz, K. (2019). Autophagy regulates steroid production by mediating cholesterol trafficking in endocrine cells. Autophagy 15, 1478–1480.

17. Kaplowitz, P.B. (2008). Link between body fat and the timing of puberty. Pediatrics 121 Suppl 3, S208–217.

18. Rodenfels, J., Lavrynenko, O., Ayciriex, S., Sampaio, J.L., Carvalho, M., Shevchenko, A., and Eaton, S. (2014). Production of systemically circulating Hedgehog by the intestine couples nutrition to growth and development. Genes Dev 28, 2636–2651.

19. Zhang, J., Liu, Y., Jiang, K., and Jia, J. (2020). Hedgehog signaling promotes lipolysis in adipose tissue through directly regulating Bmm/ATGL lipase. Dev Biol 457, 128–139.

20. Carvalho, M., Schwudke, D., Sampaio, J.L., Palm, W., Riezman, I., Dey, G., Gupta, G.D., Mayor, S., Riezman, H., Shevchenko, A., et al. (2010). Survival strategies of a sterol auxotroph. Development 137, 3675–3685.

21. Miller, W.L., and Auchus, R.J. (2011). The molecular biology, biochemistry, and physiology of human steroidogenesis and its disorders. Endocrine reviews 32, 81–151.

22. Palm, W., Sampaio, J.L., Brankatschk, M., Carvalho, M., Mahmoud, A., Shevchenko, A., and Eaton, S. (2012). Lipoproteins in Drosophila melanogaster--assembly, function, and influence on tissue lipid composition. PLoS Genet 8, e1002828.

23. Vanier, M.T. (2015). Complex lipid trafficking in Niemann-Pick disease type C. J Inherit Metab Dis 38, 187–199.

24. Castellano, B.M., Thelen, A.M., Moldavski, O., Feltes, M., van der Welle, R.E., Mydock-McGrane, L., Jiang, X., van Eijkeren, R.J., Davis, O.B., Louie, S.M., et al. (2017). Lysosomal cholesterol activates mTORC1 via an SLC38A9-Niemann-Pick C1 signaling complex. Science 355, 1306–1311.

25. Reis, T. (2016). Effects of Synthetic Diets Enriched in Specific Nutrients on Drosophila Development, Body Fat, and Lifespan. PLoS One 11, e0146758.

26. Ikeya, T., Galic, M., Belawat, P., Nairz, K., and Hafen, E. (2002). Nutrient-dependent expression of insulin-like peptides from neuroendocrine cells in the CNS contributes to growth regulation in Drosophila. Curr Biol 12, 1293–1300.

27. Rulifson, E.J., Kim, S.K., and Nusse, R. (2002). Ablation of insulin-producing neurons in flies: growth and diabetic phenotypes. Science 296, 1118–1120.

28. Walkiewicz, M.A., and Stern, M. (2009). Increased insulin/insulin growth factor signaling advances the onset of metamorphosis in Drosophila. PLoS One 4, e5072.

29. Texada, M.J., Malita, A., Christensen, C.F., Dall, K.B., Faergeman, N.J., Nagy, S., Halberg, K.A., and Rewitz, K. (2019). Autophagy-Mediated Cholesterol Trafficking Controls Steroid Production. Dev Cell 48, 659–671 e654.

30. Rewitz, K.F., Yamanaka, N., Gilbert, L.I., and O’Connor, M.B. (2009). The insect neuropeptide PTTH activates receptor tyrosine kinase torso to initiate metamorphosis. Science 326, 1403–1405.

31. Park, S., Alfa, R.W., Topper, S.M., Kim, G.E., Kockel, L., and Kim, S.K. (2014). A genetic strategy to measure circulating Drosophila insulin reveals genes regulating insulin production and secretion. PLoS Genet 10, e1004555.

32. Geminard, C., Rulifson, E.J., and Leopold, P. (2009). Remote control of insulin secretion by fat cells in Drosophila. Cell Metab 10, 199–207.

33. Koyama, T., Texada, M.J., Halberg, K.A., and Rewitz, K. (2020). Metabolism and growth adaptation to environmental conditions in Drosophila. Cell Mol Life Sci 77, 4523–4551.

34. Voght, S.P., Fluegel, M.L., Andrews, L.A., and Pallanck, L.J. (2007). Drosophila NPC1b promotes an early step in sterol absorption from the midgut epithelium. Cell Metab 5, 195–205.

35. McBrayer, Z., Ono, H., Shimell, M., Parvy, J.P., Beckstead, R.B., Warren, J.T., Thummel, C.S., Dauphin-Villemant, C., Gilbert, L.I., and O’Connor, M.B. (2007). Prothoracicotropic hormone regulates developmental timing and body size in Drosophila. Dev Cell 13, 857–871.

36. Brankatschk, M., Dunst, S., Nemetschke, L., and Eaton, S. (2014). Delivery of circulating lipoproteins to specific neurons in the Drosophila brain regulates systemic insulin signaling. Elife 3.

37. Okamoto, N., Yamanaka, N., Yagi, Y., Nishida, Y., Kataoka, H., O’Connor, M.B., and Mizoguchi, A. (2009). A fat body-derived IGF-like peptide regulates postfeeding growth in Drosophila. Dev Cell 17, 885–891.

38. Slaidina, M., Delanoue, R., Gronke, S., Partridge, L., and Leopold, P. (2009). A Drosophila insulin-like peptide promotes growth during nonfeeding states. Dev Cell 17, 874–884.

39. Layalle, S., Arquier, N., and Leopold, P. (2008). The TOR pathway couples nutrition and developmental timing in Drosophila. Dev Cell 15, 568–577.

40. Mirth, C., Truman, J.W., and Riddiford, L.M. (2005). The role of the prothoracic gland in determining critical weight for metamorphosis in Drosophila melanogaster. Curr Biol 15, 1796–1807.

41. Ohhara, Y., Kobayashi, S., and Yamanaka, N. (2017). Nutrient-Dependent Endocycling in Steroidogenic Tissue Dictates Timing of Metamorphosis in Drosophila melanogaster. PLoS Genet 13, e1006583.

42. Zeng, J., Huynh, N., Phelps, B., and King-Jones, K. (2020). Snail synchronizes endocycling in a TOR-dependent manner to coordinate entry and escape from endoreplication pausing during the Drosophila critical weight checkpoint. PLoS Biol 18, e3000609.

43. Danielsen, E.T., Moeller, M.E., Yamanaka, N., Ou, Q., Laursen, J.M., Soenderholm, C., Zhuo, R., Phelps, B., Tang, K., Zeng, J., et al. (2016). A Drosophila Genome-Wide Screen Identifies Regulators of Steroid Hormone Production and Developmental Timing. Dev Cell 37, 558–570.

44. Koyama, T., Rodrigues, M.A., Athanasiadis, A., Shingleton, A.W., and Mirth, C.K. (2014). Nutritional control of body size through FoxO-Ultraspiracle mediated ecdysone biosynthesis. Elife 3.

45. Colombani, J., Andersen, D.S., and Leopold, P. (2012). Secreted peptide Dilp8 coordinates Drosophila tissue growth with developmental timing. Science 336, 582–585.

46. Garelli, A., Gontijo, A.M., Miguela, V., Caparros, E., and Dominguez, M. (2012). Imaginal discs secrete insulin-like peptide 8 to mediate plasticity of growth and maturation. Science 336, 579–582.

47. Romao, D., Muzzopappa, M., Barrio, L., and Milan, M. (2021). The Upd3 cytokine couples inflammation to maturation defects in Drosophila. Curr Biol 31, 1780–1787 e1786.

48. Elias, C.F. (2012). Leptin action in pubertal development: recent advances and unanswered questions. Trends Endocrinol Metab 23, 9–15.

49. Krause, B.R., and Hartman, A.D. (1984). Adipose tissue and cholesterol metabolism. J Lipid Res 25, 97–110.

50. Shingleton, A.W., Das, J., Vinicius, L., and Stern, D.L. (2005). The temporal requirements for insulin signaling during development in Drosophila. PLoS Biol 3, e289.

51. Phillips, S.E., Woodruff, E.A., 3rd, Liang, P., Patten, M., and Broadie, K. (2008). Neuronal loss of Drosophila NPC1a causes cholesterol aggregation and age-progressive neurodegeneration. J Neurosci 28, 6569–6582.

52. Patel, S.C., Suresh, S., Kumar, U., Hu, C.Y., Cooney, A., Blanchette-Mackie, E.J., Neufeld, E.B., Patel, R.C., Brady, R.O., Patel, Y.C., et al. (1999). Localization of Niemann-Pick C1 protein in astrocytes: implications for neuronal degeneration in Niemann-Pick type C disease. Proc Natl Acad Sci U S A 96, 1657–1662.

53. Bambace, C., Dahlman, I., Arner, P., and Kulyte, A. (2013). NPC1 in human white adipose tissue and obesity. BMC Endocr Disord 13, 5.

54. Meyre, D., Delplanque, J., Chevre, J.C., Lecoeur, C., Lobbens, S., Gallina, S., Durand, E., Vatin, V., Degraeve, F., Proenca, C., et al. (2009). Genome-wide association study for early-onset and morbid adult obesity identifies three new risk loci in European populations. Nat Genet 41, 157–159.

55. Al-Daghri, N.M., Cagliani, R., Forni, D., Alokail, M.S., Pozzoli, U., Alkharfy, K.M., Sabico, S., Clerici, M., and Sironi, M. (2012). Mammalian NPC1 genes may undergo positive selection and human polymorphisms associate with type 2 diabetes. BMC Med 10, 140.

56. Mariman, E.C., Bouwman, F.G., Aller, E.E., van Baak, M.A., and Wang, P. (2015). Extreme obesity is associated with variation in genes related to the circadian rhythm of food intake and hypothalamic signaling. Physiol Genomics 47, 225–231.

57. Liu, R., Zou, Y., Hong, J., Cao, M., Cui, B., Zhang, H., Chen, M., Shi, J., Ning, T., Zhao, S., et al. (2017). Rare Loss-of-Function Variants in NPC1 Predispose to Human Obesity. Diabetes 66, 935–947.

58. Uronen, R.L., Lundmark, P., Orho-Melander, M., Jauhiainen, M., Larsson, K., Siegbahn, A., Wallentin, L., Zethelius, B., Melander, O., Syvanen, A.C., et al. (2010). Niemann-Pick C1 modulates hepatic triglyceride metabolism and its genetic variation contributes to serum triglyceride levels. Arterioscler Thromb Vasc Biol 30, 1614–1620.

59. Kuzu, O.F., Noory, M.A., and Robertson, G.P. (2016). The Role of Cholesterol in Cancer. Cancer Res 76, 2063–2070.

60. Stork, T., Engelen, D., Krudewig, A., Silies, M., Bainton, R.J., and Klambt, C. (2008). Organization and function of the blood-brain barrier in Drosophila. J Neurosci 28, 587–597.

61. Jenett, A., Rubin, G.M., Ngo, T.T., Shepherd, D., Murphy, C., Dionne, H., Pfeiffer, B.D., Cavallaro, A., Hall, D., Jeter, J., et al. (2012). A GAL4-driver line resource for Drosophila neurobiology. Cell Rep 2, 991–1001.

62. Masuyama, K., Zhang, Y., Rao, Y., and Wang, J.W. (2012). Mapping neural circuits with activity-dependent nuclear import of a transcription factor. J Neurogenet 26, 89–102.

63. Ono, H., Rewitz, K.F., Shinoda, T., Itoyama, K., Petryk, A., Rybczynski, R., Jarcho, M., Warren, J.T., Marques, G., Shimell, M.J., et al. (2006). Spook and Spookier code for stage-specific components of the ecdysone biosynthetic pathway in Diptera. Dev Biol 298, 555–570.

64. Pan, X., Neufeld, T.P., and O’Connor, M.B. (2019). A Tissue- and Temporal-Specific Autophagic Switch Controls Drosophila Pre-metamorphic Nutritional Checkpoints. Curr Biol 29, 2840–2851 e2844.

65. Bader, R., Sarraf-Zadeh, L., Peters, M., Moderau, N., Stocker, H., Kohler, K., Pankratz, M.J., and Hafen, E. (2013). The IGFBP7 homolog Imp-L2 promotes insulin signaling in distinct neurons of the Drosophila brain. J Cell Sci 126, 2571–2576.

66. Veenstra, J.A. (2009). Peptidergic paracrine and endocrine cells in the midgut of the fruit fly maggot. Cell Tissue Res 336, 309–323.

67. Yamanaka, N., Romero, N.M., Martin, F.A., Rewitz, K.F., Sun, M., O’Connor, M.B., and Leopold, P. (2013). Neuroendocrine control of Drosophila larval light preference. Science 341, 1113–1116.

68. Li, Q., and Gong, Z. (2015). Cold-sensing regulates Drosophila growth through insulin-producing cells. Nat Commun 6, 10083.

69. Varela, N., Gaspar, M., Dias, S., and Vasconcelos, M.L. (2019). Avoidance response to CO2 in the lateral horn. PLoS Biol 17, e2006749.

