## Supplemental table 1: qPCR oligo sequences for "Insulin signaling couples growth and early maturation to cholesterol intake"

| Oligo name | Source of sequence |
|---|---|
| *4EBP* -F: CCAGGAAGGTTGTCATCTCG | Moeller *et al.* , 2017 |
| *4EBP* -R: CCAGGAGTGGTGGAGTAGAGG | Moeller *et al.* , 2017 |
| *CCHa2* -F: GCCTACGGTCATGTGTGCTAC | Sano *et al.* , 2015 |
| *CCHa2* -R: ATCATGGGCAGTAGGCCATT | Sano *et al.* , 2015 |
| *E75a* -f: ACCACAGCACCACCCATTT | Christensen *et al.,* 2020 |
| *E75a* -r: TGTTTGGCGGTAGTTTCAGG | Christensen *et al.,* 2020 |
| *E75b* -f: CAACAGCAACAACACCCAGA | Christensen *et al.,* 2020 |
| *E75b* -r: CAGATCGGCACATGGCTTT | Christensen *et al.,* 2020 |
| *Gbp1* -F: TGTGCTGCCAGTTATAGACAAC | FlyPrimerBank |
| *Gbp1* -R: TCGTTTTCCGAACAGCTCAAA | FlyPrimerBank |
| *Gbp2* -F: AGGTTGGCTATGTCACTGATCC | FlyPrimerBank |
| *Gbp2* -R: ACCCATGTACGTGATGACCAT | FlyPrimerBank |
| *Ilp2* -F: CTCAACGAGGTGCTGAGTATG | Texada *et al.,* 2019 |
| *Ilp2* -R: GAGTTATCCTCCTCCTCGAACT | Texada *et al.,* 2019 |
| *Ilp3* -F: CAACGCAATGACCAAGAGAAC | Texada *et al.,* 2019 |
| *Ilp3* -R: GCATCTGAACCGAACTATCACTC | Texada *et al.,* 2019 |
| *Ilp5* -F: ATGGACATGCTGAGGGTTG | Texada *et al.,* 2019 |
| *Ilp5* -R: GTGGTGAGATTCGGAGCTATC | Texada *et al.,* 2019 |
| *Ilp6* -F: TGCTAGTCCTGGCCACCTTGTTCG | Okamoto *et al* ., 2009 |
| *Ilp6* -R: GGAAATACATCGCCAAGGGCCACC | Okamoto *et al* ., 2009 |
| *InR* -F: CTCAGCCATACCAGGGACTTT | Moeller *et al.* , 2017 |
| *InR* -R: CTCTCCATAACACCGCCATC | Moeller *et al.* , 2017 |
| *Ptth* -F: TGAAGGTTTGCACGAGATGG | Christensen *et al.,* 2020 |
| *Ptth* -R: CTGTGGGATGTGGAGTGCTG | Christensen *et al.,* 2020 |
| *Rp49* -F: AGTATCTGATGCCCAACATCG | Sano *et al.* , 2015 |
| *Rp49* -R: CAATCTCCTTGCGCTTCTTG | Sano *et al.* , 2015 |
| *sun* -F: ATGACTGCCTGGAGAGCTG | FlyPrimerBank |
| *sun* -R: GTGAACTTCACATGGCTCGC | FlyPrimerBank |
| *upd2* -F: CGGAACATCACGATGAGCGAAT | Rajan and Perrimon, 2012 |
| *upd2* -R: TCGGCAGGAACTTGTACTCG | Rajan and Perrimon, 2012 |
